# Modeling viral and bacterial infections in human lung organotypic systems reveals strain specific host responses

**DOI:** 10.1101/2025.03.31.644992

**Authors:** Bárbara Faria Fonseca, Jérôme Wong-Ng, Michael Connor, Héloïse Mary, Min Hee Kim, Rémy Yim, Vincent Bondet, Vincent Michel, Hélène Strick Marchand, James Di Santo, Darragh Duffy, Mélanie Hamon, Nathalie Sauvonnet, Lisa A. Chakrabarti, Samy Gobaa

## Abstract

In this study, we developed novel lung organoid-on-chip models that elucidate differential human tissue response to various strains of respiratory pathogens: *Streptococcus pneumoniae* and SARS-CoV-2. We show that human fetal-derived distal lung epithelial cells are readily expandable in 3D as organoids, thereby providing a highly sustainable source of lung progenitor cells. These 3D organoid progenitors can then be induced to produce airway and alveolar organoids on microfluidic devices. Upon challenge with *Streptococcus pneumoniae*, a bacterium known to cause pneumonia, a rapid and strain-dependent colonization was observed at the epithelial surface of alveolar chips. We also assessed SARS-CoV-2 infection in the alveoli-on-chip system and observed that the Delta variant exhibited greater infectivity as compared to the Omicron BA.5. Both SARS-CoV-2 variants induced potent interferon responses and triggered the expression of different interferon-stimulated genes. Our results demonstrate that strain-specific host defense mechanisms can be recapitulated in human-organoid-based microfluidic systems, paving the way for the use of such platforms for more targeted assessments of human response to novel emergent pathogen strains.

**Highlights:**

- Human fetal epithelial lung stem cells can be expanded as multipotent organoids and differentiated into both airway or alveolar organoids
- Multipotent lung organoids efficiently produce functional epithelia of small airway or alveoli when grown on-chip.
- *Streptococcus pneumoniae* inoculation in alveoli-on-chip mimics the early stages of bacterial colonization in lung epithelia
- Alveoli on-chip system recapitulates variant-specific interactions. SARS-CoV-2 Delta replicates but not Omicron BA.5.
- Robust interferon response upon SARS-CoV-2 infection shows Alveoli on-chip can model innate immune responses.

## Introduction

The epithelial lining of the human respiratory tract constitutes the major entry point for airborne pathogens. At the tissue level, lung epithelia are supported by a highly specialized stroma, a richly vascularized network and numerous populations of specialized immune cells. Collectively, this complex system ensures tissue homeostasis, barrier function and efficient gas exchange while facing constant viral and bacterial challenges. Host-microbe interactions start at the epithelial apical surface. Resulting infections trigger early defense mechanisms including mucociliary clearance and mucosal innate immunity ^1,2^. The complexity of the mounted responses is in part determined by the heterogeneous histological composition of the tissues along the respiratory tract. Ciliated and mucus-producing cells in the airways work together to remove airborne pollutants and microbes. At the distal end of the tract, the alveolar epithelium can also mount a complex immune response in response to pathogen invasion. Disentangling the lung tissue response to airborne pathogens hinges on the establishment of advanced tissue models that extend beyond what can be recapitulated in standard 2D, cell line-based assays.

The major challenge in devising advanced human lung disease models resides in establishing a sustainable source of human cells. Ideally, the starting cell material should be amenable to *in vitro* expansion while still capable of producing the complete range of cell types that constitute the tissue of interest. Immortalized cell lines, such as A549 and Calu-3, can be useful for drug screening and infection studies, but as they originate from lung carcinoma, they poorly represent the biology of bona fide lung cells ^3^. On the other hand, primary adult cells can be used to reconstitute nasopharyngeal, bronchial and alveolar tissues *in vitro* ^4–6^. However, these largely postmitotic cells can be difficult to obtain and rapidly dedifferentiate ^7^. To overcome these limitations, culturing primary lung cells as organoids has proven highly effective. Both human bronchiolar and alveolar organoids can today be derived from human adult biopsies or from iPSC cell culture ^8–15^. Lung organoids can be maintained for several passages and be used for lung disease modeling ^16,17^.

Reproducing the apicobasal polarity of respiratory epithelia is crucial for correctly modeling the entry of an airborne pathogen in a human lung cell. Most current lung organoid protocols lead to the production of apical surfaces facing inward (lumen). Loading any inoculum on the apical side can therefore be challenging. Various strategies were employed to partially overcome this limitation including microinjection ^18,19^, inverted-polarity techniques ^10^ and organoid fragmentation and re-formation ^20^. However, these approaches often compromise throughput and reproducibility. Alternatively, organ-on-chip technology was shown to be a powerful technique to reconstruct accessible human epithelial tissues in an environment where tissue perfusion of media, drugs and pathogens can be precisely controlled ^4,21,22^. This microfluidic-based approach also enables the creation of interconnected channels and can allow the stretching of cultured epithelia ^4,23^. Additionally, organ-on-chip systems allow human lung epithelial cells to be cultured at an air-liquid interface (ALI), further mimicking the function of the human respiratory tract. Leveraging these technologies for investing host-pathogen interactions is gaining traction. However, new developments remain pathogen-dependent.

In this study, we combined two organotypic models, organoids and organ-on-chips, to build novel small airway and alveolar lung infection models. We expanded human distal lung bud organoids ^24–26^ and steered them to differentiate, on-chip, into sample-matched airway or alveolar tissues. We demonstrate that such advanced organ-on-chip devices can successfully model viral and bacterial infections. We found that the engineered lung models can be used to identify cell type-related susceptibilities to bacterial infection. We also show that the presented approaches can be used to characterize variant-dependent human innate immune responses upon SARS-CoV-2 infection.

## Results

### Derivation of human multipotent lung organoids

During human lung development, the epithelial lung bud tip forms a pool of multipotent stem cells capable of generating both small airways and alveoli tissues. A complex interplay of signaling pathways, primarily driven by WNT and fibroblast growth factor (FGF) activation, ensure the plasticity of this fetal structure along the pseudoglandular and early-canalicular stages of lung development ^2^. In vitro, reconstructing this signaling cocktail allows long-term expansion of lung multipotent progenitors ^24^. In this study, we derived a stable culture of human fetal-derived lung bud tip organoids that can be amplified and differentiated into either alveolar or small airway epithelial cells (Figure 1A). Lung organoids cultured in self-renewing conditions grow as simple epithelia that are SOX2+ and SOX9+ (Figure 1B & 1D) ^2^. These organoids could be split every 10 to 14 days, multiple times, to expand the number of available cells (Figure 1C). We also found that stable multipotent organoid cultures can be successfully reestablished after multiple freeze-thaw cycles, enabling the banking of this initial material, which greatly simplifies its use and dissemination.

**Figure 1.**
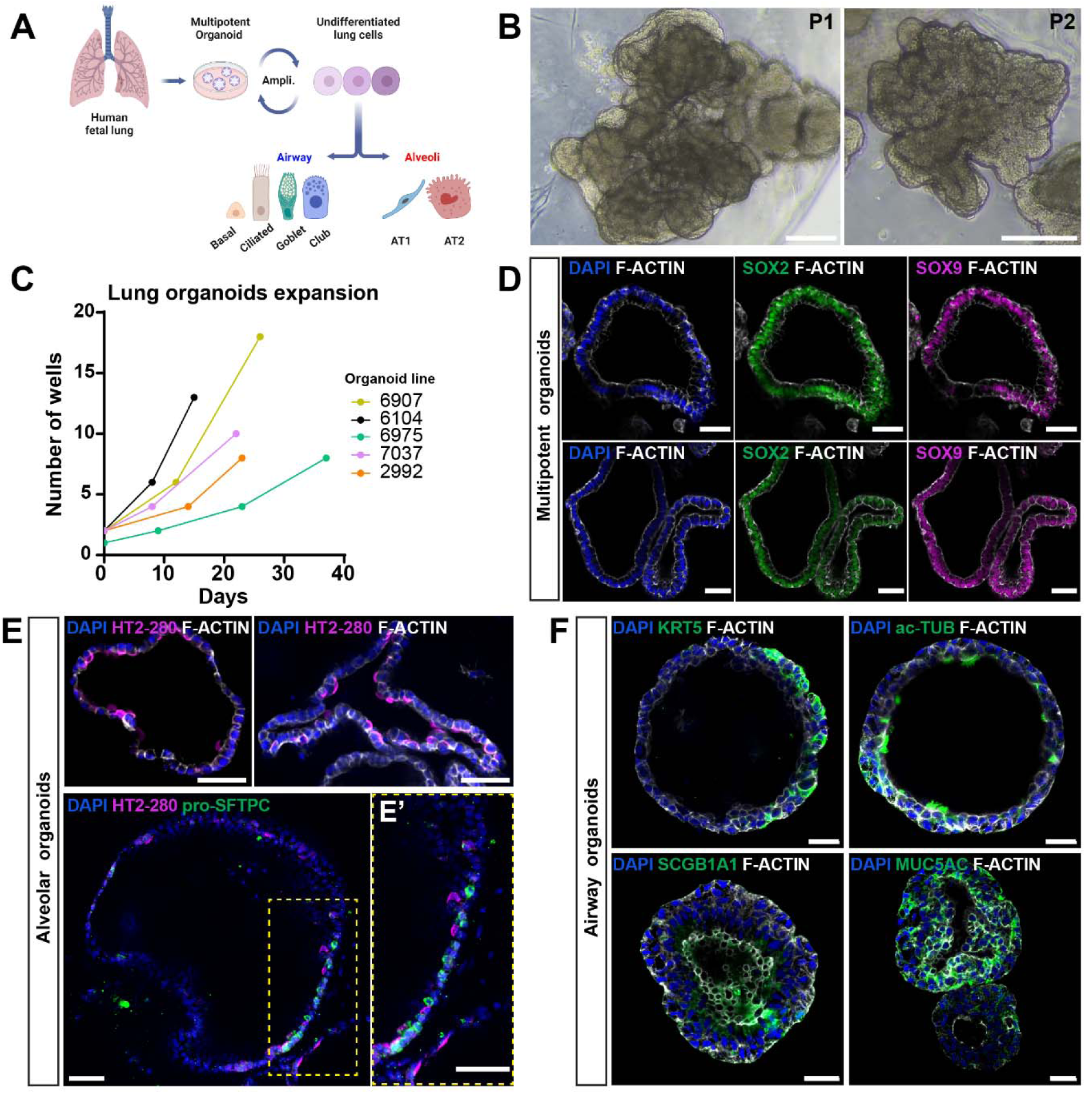
Derivation of three types of human fetal lung organoids for the production of airway and alveolar cell types. (A) Schematic representation of the cell culture pipeline used to produce different fetal lung organoid types. (B) Brightfield images of multipotent lung organoids in self-renewing conditions at passage 1 (left) and passage 2 (right). (C) Expansion of the organoid cultures over multiple passages. Each line corresponds to one experiment with a distinct organoid line. (D) Multipotent lung organoids showing positive immunostainings for SOX2 and SOX9 markers (E) Alveolar organoids showing typical alveolar type 2 cells identified with HT2-280 and anti-pro-SFTPC immunolabeling. (E’) Higher magnification of the region depicted in (E). (F) Immunostaining of airway organoids showing typical cell types including basal (KRT5+), ciliated (ac-TUB+), club (SCGB1a1+) and goblet (MUC5AC+) cells. Scale bars: 75 µm (B), 50 µm (D and E), 20 µm (F).

To steer lung multipotent organoids towards alveolar differentiation, we modified the culture conditions by withdrawing FGF, maintaining WNT activation and adding NOTCH and TGF-β inhibitors, as described in Lim et al ^26^. After two weeks in these differentiation conditions, immunostaining revealed the presence of HT2-280 positive cells, a human lung-specific type II pneumocytes (AT2) marker ^27^. Pro-surfactant protein C (pro-SFTPC), a typical surfactant protein produced by bipotent alveolar progenitors and fully differentiated AT2 cells ^2^, was also identified in several cells. These observations confirm the differentiation into AT2 cells (Figure 1E). Here, we did not identify cells with positive labeling for the HT1-56, AQP5 and RAGE, markers of type I pneumocytes (ATI). While extensive differentiation dynamics were not investigated, we speculate that terminal AT1 differentiation within organoids requires additional extrinsic factors including morphogens or physical cues to support their development. Previous studies with alveolar organoids derived from primary adult lung cells also highlighted an absence of AT1 cells ^8,28^. Interestingly, recent reports with iPSC-derived and fetal-derived alveolar organoids show that withdrawal of WNT agonists, combined with YAP activators in the differentiation medium, is necessary for AT1 differentiation ^14,29^.

Next, we sought to evaluate the capacity of multipotent organoids to differentiate into small airway tissues, lung organoids were treated with a pro-airway differentiation medium where WNT agonists were removed and FGF activation maintained ^9^. A two-week treatment was sufficient to induce the differentiation into typical airway cell types such as beating multiciliated cells that are positive for acetylated tubulin (ac-TUB+) and basal cells expressing cytokeratin 5 (KRT5+). We also identified secretory cells, including goblet cells producing mucin (MUC5AC+) and club cells expressing secretoglobin family 1A member 1 (SCGB1A1+) (Figure 1F; Supplementary Movie 1). Our observations show that the morphology of differentiated organoids showed varying lumen sizes and different extents of folding but systematically appeared as pseudostratified when compared to multipotent organoids.

Altogether, we demonstrated that human fetal distal lung buds can be efficiently harvested and used as a reliable source material for the derivation of multipotent human lung organoids. We also showed that these organoids have the capacity to differentiate into either small airways or alveolar epithelia.

### On-chip differentiation of lung reconstituted epithelia

Devising human lung organotypic models to investigate infectious diseases relies first on the reconstitution of faithful mimics of human epithelia and second on straightforward access to the luminal compartment that is the classical entry point of airborne pathogens. While organoids can efficiently ensure an appropriate cell composition, their inward polarization renders infectious studies labor intensive ^18,19^. Here we engineered an organ-on-chip approach that relies on the use of human organoids as a source of cells and where both the perfusion of fluids and the inoculum loading are fully controlled ^4,21,22^.

Briefly, we seeded fragments of multipotent organoids on a commercial S1™ organ-on-chip device from emulate™ (Figure 2A). Two days of culture under self-renewing conditions were sufficient to produce a confluent epithelium on the chips. We also observed that adding sample-matched mesenchymal cells to the lower channel of the chip drastically improved lung epithelial cell survival and differentiation as previously reported ^6,8,26,30,31^. Next, we induced differentiation by perfusing either airway or alveolar media optimized for organoid differentiation. Finally, air-liquid interface (ALI) was implemented to promote differentiation (Figure 2A). Both types of organoid-derived chips formed a simple epithelium expressing E-cadherin and ZO-1 proteins (Supplementary Figure 1). The produced epithelia maintained ALI for multiple days, suggesting that the junctions are tight.

**Figure 2:**
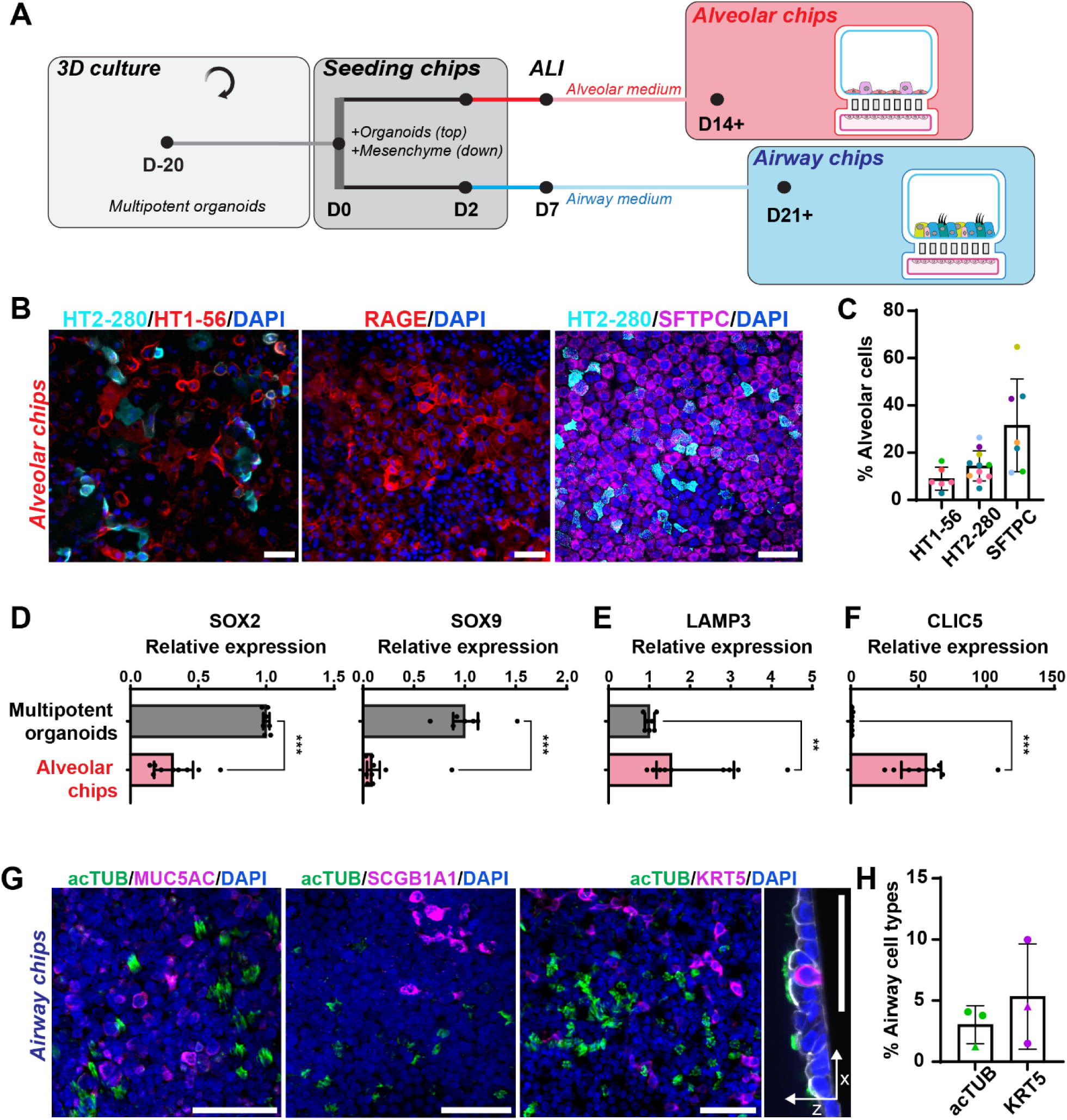
Characterization of human airway-on-chip and alveoli-on-chip systems. (A) Schematic representation of the protocol used to derive alveolar and airway organoid-on-chip devices. (B) Immunolabelling of the alveoli-on-chip system. Left: AT1 (HT1-56+), AT2 (HT2-280+); middle: AT1 (RAGE+); right: AT2 (HT2-280+, pro-SFTPC+). (C) Percentage of cells labelled with the following alveolar markers: HT1-56, HT2-280 and pro-SFTPC. Each dot color corresponds to a different organoid line. Mean +/- standard deviation (SD) are plotted. (D-F) Comparative RT-qPCR analyses between multipotent organoids versus lung cells in the alveolar chips showing the relative expression of typical lung multipotent markers (SOX2 and SOX9) (D), the mature AT2 marker LAMP3 (E) and the mature AT1 marker CLIC5 (F) (D-F: Median +/- interquartile range are plotted and statistically significant differences are assessed with the Mann-Whitney test. n=4 independent experiments with 4 different samples). (G) Immunolabelling of airway cell types in the airway-on-chip system. Left: ciliated (ac-TUB+) and goblet (MUC5AC+) cells; middle: ciliated (ac-TUB+) and club (SCGB1A1+) cells; right: ciliated (ac-TUB+) and basal (KRT5+) cells. (H) Percentage of ciliated (ac-TUB+) and basal (KRT5+) cells in airway chips. Mean +/- SDs are plotted. (B, G) Scale bars: 50µm.

Next, we analyzed the cell composition of chips induced to differentiate toward the alveolar lineage (Figure 2B-F). Immunostaining analysis revealed that approximately one third of the cells expressed pro-SFTPC (31.6% ±19.6), (Figure 2B, 2C). HT2-280, stained an average of 14.4% ±6.4 of cells, indicating that only 12 days in differentiation medium was sufficient to drive the differentiation of a substantial population of AT2 cells on-chip. Interestingly, AT1 cells could be identified with RAGE and HT1-56 antibodies, the latter labelling 9.0 ± 4.8% of the cells (Figure 2B, 2C). RT-qPCR analyses confirmed a down-regulation in the expression of bud tip markers (SOX2 and SOX9, Figure 2D) and an up-regulation on the expression of mature alveolar markers including LAMP3 (a member of the lysosomal-associated membrane protein family and marker of mature AT2 cells ^32^) and chloride intracellular channel 5 (CLIC5), expressed by bona fide AT1 cells ^33^ (Figure 2E, 2F). Finally, we checked for incidental airway differentiation on alveolar chips and found no evidence of the presence of ciliated, goblet or club cells presence. Basal cells, positive for cytokeratin 5 (KRT5) and transcription factor SOX2 but negative for SOX9, were very seldomly observed. We termed these devices AlveoChips.

When analyzing the cellular composition of small airway chips, immunostaining showed a standard profile of cells including ciliated cells (acTUB+), club cells (SCGB1A1+), goblet cells (MUC5AC+) and basal cells (KRT5+) (Figure 2G, 2H). In addition, a clearly polarized, pseudo-stratified epithelium was observed (Figure 2G). Time-lapse microscopy also showed the presence of beating cilia at the surface of the reconstituted airway epithelium (Supplementary Movie 2) after 20 days of differentiation. Basal cells accounted for 5.3% ± 4.3 and multi-ciliated cells for 3%± 1.6 of the total cells in the epithelium (Figure 2H). These data show that, as for the organoid cultures, chips seeded with human fetal cells derived from multipotent organoids can be guided to produce either airway or alveolar microphysiological systems.

### AlveoChip models *Streptococcus pneumoniae* alveolar colonization

*S. pneumoniae* colonizes the human upper respiratory tract where it is carried asymptomatically. However, when the bacterium invades the alveolar sacs, it can cause severe pneumoniae ^34^. Our understanding of the infectious processes, at early stages of pneumococcal colonization, that drive asymptomatic carriage versus pathogenic invasion is limited. Therefore, studying the early events of *S. pneumoniae* in reconstituted human primary lung tissue is highly relevant. Due to the accurate epithelial cell composition of AlveoChips and the capacity to finely control the inoculum administration and subsequent infectious events through microfluidics, we aimed to model *S. pneumoniae* colonization on the alveolar epithelium. Here we used two *S. pneumoniae* strains, TIGR4 and 6B ST90, which are known to cause disease in human and in murine models, with TIGR4 being more invasive than 6B ST90 ^35^. For comparison, we used *Corynebacterium accolens* (*C. accolens*), a commensal nasal microbiont that does not colonize the lungs ^36^.

Briefly, we inoculated AlveoChips TIGR4, 6B ST90 or *C.accolens* bacteria for 6 min under flow (1000ul/h). We then removed all the medium in the top channel and re-established an air-liquid interface (Figure 3A). A dose of 3.0x 10^4^ colony-forming units (CFU) was used on every AlveoChip (Figure 3B). At 24 hours post infection (hpi), we observed an approximate 1.5 order of magnitude increase of CFU counts in top channel effluents collected from both TIGR4 (median: 5.0.10^5^ total CFU) and 6B ST90 infected chips (median: 7.9.10^5^ total CFU) (Figure 3C). Interestingly, a 2-log decrease in the number of CFU counts was observed in effluents of chips treated with *C. accolens* (median: 3.3.10^2^ total CFU). These results show that the two pneumococcus strains can successfully multiply in the modeled alveolar microenvironment. In contrast, *C. accolens* does not replicate and its inoculated population declines rapidly within the first 24hpi.

**Figure 3:**
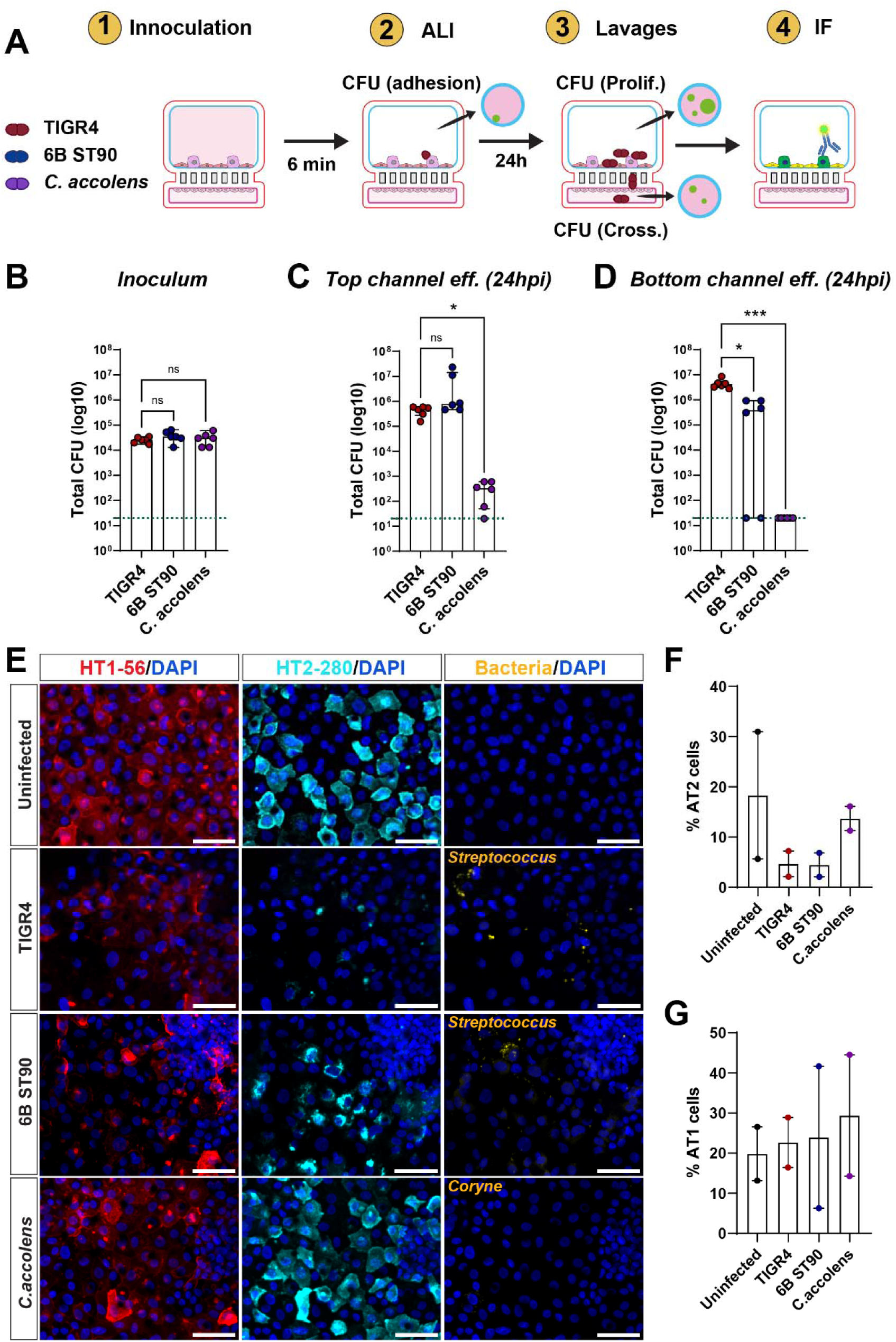
*Streptococcus pneumoniae* strains infect the human alveoli on-chip system. (A) Schematic representation of the infection protocol with *S. pneumoniae* strains (TIGR4 and 6B ST90) or *C. accolens* (commensal control). ALI: air-liquid-interface, IF: immunofluorescence (B) Colony forming unit (CFU) counts the bacteria inoculum. (C) CFU counts of respective bacteria collected after 24h in the top channel effluent. (D) CFU counts of respective bacteria collected after 24h in the bottom channel effluent. Median +/- interquartile range are plotted and statistically significant differences between bacteria were compared with Kruskal-Wallis test. (n=3 independent experiments with 3 different organoid lines; geometric means of 2 technical counts per chip are shown). Dotted green in lines B-D: limit of detection value (20 total CFU). (E) Representative images of bacterium infection in alveolar chips. HT1-56 and HT2-280 immunolabeling identifies AT1 and AT2 cells, respectively. hpi: hours post-infection. Scale bars: 50µm. (F and G) Percentages of AT2 and AT1 cells upon bacterial infections on-chip. Mean +/- standard deviation, n=2 independent experiments. Minimum cell count per experiment above 20000.

We next asked whether these microorganisms could be found in effluents of the bottom channel, suggesting a possible transmigration towards the mesenchyme-filled compartment. High levels of TIGR4 pneumococcus counts were detected at the bottom channel effluents (median: 4.1.10^6^ total CFU). Interestingly, while the commensal-like strain 6B ST90 could also be detected, it was recovered at lower levels than that of TIGR4 (median: 3.9.10^5^ total CFU). CFU counts from *C. accolens* treated chips were below the limit of detection, confirming the absence of growth of this pathogen in alveolar chips (Figure 3D). CFU assays revealed that both pneumococcus strains, which have contrasted effects in mice ^35^, were able to proliferate in the AlveoChips.

Finally, immunolabeling after 24hpi revealed the presence of numerous TIGR4 and 6B ST90 bacteria but no *C. accolens*, at the surface of the epithelial cells corroborating the CFU results (Figure 3E). We could also detect TIGR4 and 6B ST90 within the pores of the chip membrane, suggesting a transmigration across the barrier to the bottom chamber (Supplementary Figure 2A). Interestingly when assessing the frequency of each cell type in infected chips, we measured a strong decrease in HT2-280+ cell occurrence upon pneumococcal invasion (Figure 3F). This decrease seemed specific to type II pneumocytes as no variation in ATI or in total cell numbers were observed after TIGR4 or 6B ST90 inoculation (Figure 3G, Supplementary Figure 2B). Our data demonstrate that *S. pneumococcus* is readily capable of colonizing the AlveoChip model and of crossing the epithelial barrier within the first 24hpi.

### AlveoChip unveils distinct replicative fitness of SARS-CoV-2 variants

To investigate the replication competence of SARS-CoV-2 on-chip, we infected AlveoChips with the SARS-CoV-2 Delta variant. Chips where then analyzed at 4 days post infection (4dpi) (Figure 4A). Immunostaining experiments detected 1.9 ± 5.5 % of SARS-CoV-2 spike+ cells (Figure 4B and 4C). Quantifications showed that 34.1 ± 27 % of the HT2-280+ were positive for the spike viral protein (Figure 4D). Interestingly, in our experimental conditions, Delta infection did not trigger major cell death as no significant differences were observed in the total number of HT2-280+ cells or in the overall cell count on chip post-infection (Supplementary Figure 3A and 3B).

**Figure 4:**
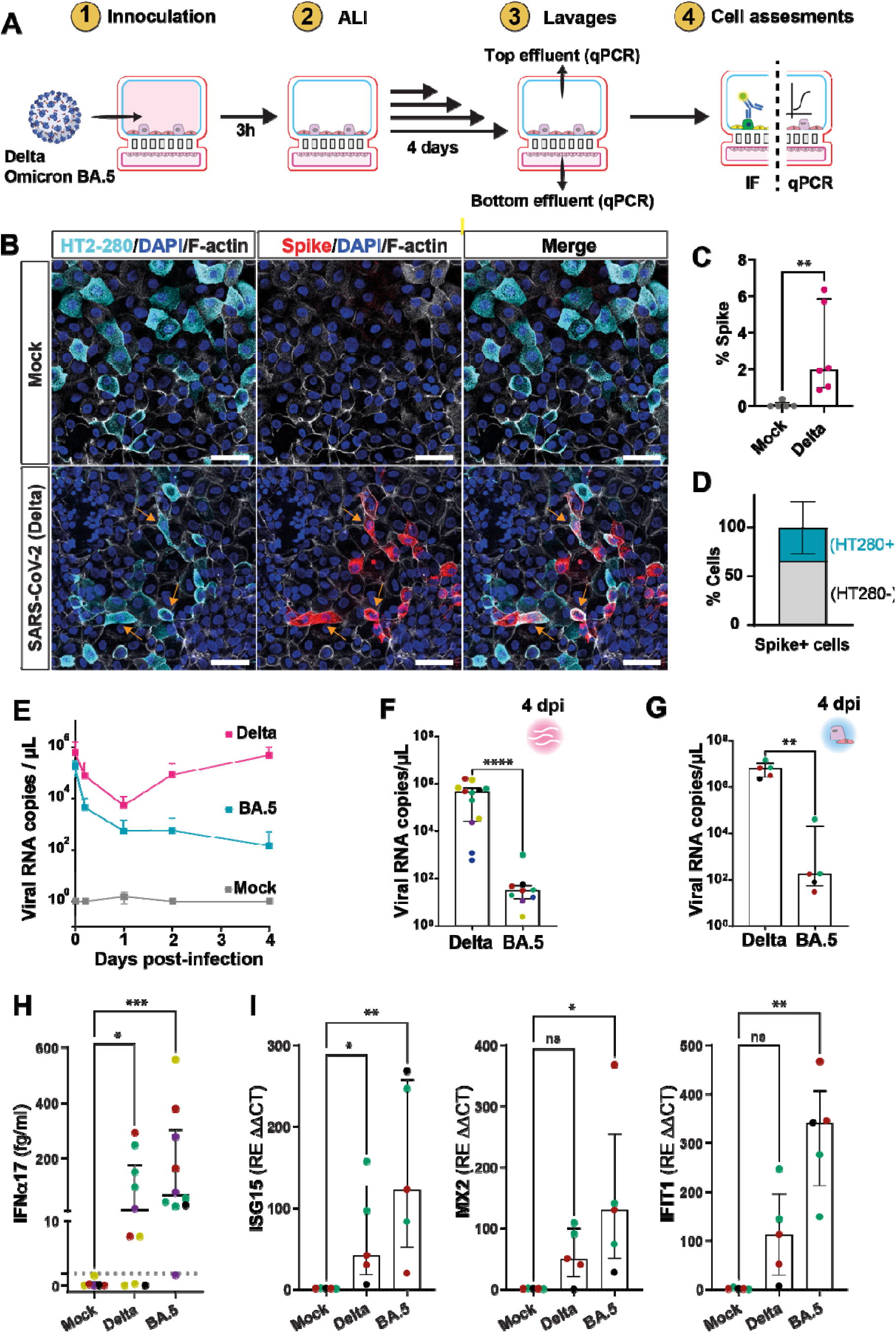
SARS-CoV-2 variant Delta, but not Omicron BA.5, replicates in human alveolar chips. (A) Schematic representation of SARS-CoV-2 infection protocol in alveoli-on-chip system. ALI: air-liquid-interface (B) Detection of SARS-CoV-2 (Delta) infected cells by immunolabeling with anti-spike antibodies (red staining). HT2-280 antibody (in cyan) stains AT2 cells. Counterstaining with DAPI and F-actin (phalloidin, in gray). Orange arrows: spike and HT2-280 co-labelled cells. Scale bars: 50 µm. (C) Percentage of spike+ cells in Delta versus mock-infected conditions. Median +/- interquartile range (IQR) are plotted and statistically significant differences are compared with the Mann-Whitney test. (n= 4 independent experiments with 3 organoid lines). (D) Proportion of HT2-280+ cells among spike+ cells. Mean +/- standard deviation (SD) are plotted. (E) Kinetics of SARS-CoV-2 viral load in top channel effluents measured by RT-qPCR. Mean +/- SD are plotted. (F) Viral RNA load in top channel effluent at 4 dpi. Median +/- IQR are plotted and statistically significant differences are compared with the Mann-Whitney test. Each dot color represents a different organoid line. (E-F, n= 6 independent experiments with 6 organoid lines). (G) Viral RNA detection in cellular extracts at 4 dpi. Median +/- IQR are plotted and statistically significant differences are compared with the Mann-Whitney test. Each dot color represents a different organoid line. (n= 3 independent experiments with 3 organoid lines). (H) IFNα protein concentrations, expressed as IFNα17 equivalents, detected in top channel washes recovered at 4 dpi (n= 5 independent experiments with 5 organoid lines. Each dot color represents an organoid line. Median +/- interquartile range are plotted and Kruskal-Wallis tests are used for addressing statistically significant differences). (I) Relative RNA expression of ISG15, MX2 and IFIT1 at 4 dpi (n=3 independent experiments with 3 organoid lines Median +/- IQR are plotted and statistically significant differences are compared with the Kruskal-Wallis test. Each dot color represents a different organoid line).

To compare the fitness of different SARS-CoV-2 variants, we infected AlveoChips with equal titers of SARS-CoV-2 Delta or Omicron BA.5 variants and then quantified the apical release of viral RNA over 4 days (Figure 4A). RT-qPCR data showed a steady replication of the Delta variant that starts at 1dpi and reaches a peak around 4dpi (Figure 4E). In contrast, Omicron BA.5 did not replicate efficiently in our alveolar model: the corresponding viral RNA load, measured in chip lavages, remained at significantly lower levels (Figure 4E). At the end of the experiment, the apically released viral RNA for the two variants differed by four orders of magnitude (4.4.10^5^ copies per mL for Delta versus 3.10^1^ copies per mL for Omicron BA.5). Interestingly this significant difference (p < 0.0001) was consistently observed across all the used organoid lines, demonstrating the high reproducibility of the system (Figure 4F). At the end of the experiment, a higher amount of viral RNA could also be detected in total cell RNA extracted from Delta-infected chips (Figure 4G). Viral particles could be identified by scanning electron microscopy in chips infected with Delta, but not in chips in mock-infected samples (Supplementary Figure 4). In addition, immunostainings performed with an anti-viral nucleocapsid (NCP1) antibody, further confirmed that SARS-CoV-2 infected cells could be detected in Delta-infected chips, but not in Omicron BA.5 and mock-infected chips (Supplementary Figure 5). Taken together, these results show that the AlveoChip model is susceptible to Delta infection, but not to Omicron BA.5, recapitulating the lower replicative potential of Omicron in an alveolar environment ^37,38^. To our knowledge, Alveochip is the first in vitro alveolar organ-on-chip system that recapitulates differential variant sensitivity.

Finaly, following viral infection, we quantified type I and III interferon (IFN) release on the luminal surface of lung epithelial cells to verify whether the reconstituted alveolar epithelia can mount an innate host-defense response. A significant increase in IFNα protein levels (expressed as IFNa17 equivalents) was observed at 4dpi in Delta-infected chips, compared to uninfected controls. Interestingly, despite the lower infectivity of Omicron BA.5 in this system, IFNα secretion was similarly high in epithelial cells challenged with this variant (Figure 4H). For IFNβ, average protein levels were higher in Omicron BA.5-infected chips, while no significant IFNλ3 production was detected at 4dpi for either variant (Supplementary Figure 6A and 6B). When looking at the transcription of interferon-stimulated genes (ISGs), both Delta and Omicron BA.5 infections triggered the upregulation of standard ISG such as IFIT1, ISG15 and MX2 (Figure 4I). Overall, these data show that, in our AlveoChip model, host epithelial cells respond to SARS-CoV-2 viral inoculation by secreting IFN molecules and expressing ISGs, thereby initiating an innate immune response. Interestingly, despite the lower replication competence of Omicron BA.5 compared to the Delta, the former still triggers a strong IFN response.

## Discussion

In this work, we assessed the potential of human fetal-derived lung organoids as starting material for building respiratory infection models. We showed that multipotent fetal lung organoids can be largely expanded *in vitro* while preserving their capacity to differentiate into small airway or alveolar epithelia. Using these lung organoids, we generated airway- and alveoli-on-chip systems which grant easy access to the apical side of epithelial cells and thus can be readily used for addressing airborne pathogen infections in well-controlled conditions.

Culturing primary adult lung cells as organoids has been extensively explored in the past decade ^8–10,39^. Both biochemical and mechanical cues provided by the extracellular matrix facilitate the self-organization and differentiation of stem cells and progenitors into faithful tissue mimics. However, several limitations, including dedifferentiation, limited supply, and donor-to-donor variations, remain to be solved. While fetal lung tissue is undeniably in very short supply, fetal lung bud tips can be expanded *in vitro* to form self-renewing organoids that preserve key markers of multipotent progenitors ^24^. On the other hand, induced pluripotent stem cells (iPSCs) can also generate lung organoids for infection studies ^40,41^. However, because it is necessary to recapitulate all the differentiation steps to reach the lung progenitor state, iPSC protocols tend to be longer and more labor-intensive than fetal lung cultures ^11–15^. Consequently, our approach provides a valuable addition to existing lung-on-chip models ^4,23,42–45^, as our system recapitulates lung epithelia composition *in vitro* without constantly relying on primary cell sourcing or on less physiological immortalized cell lines.

Airborne pathogens typically infect through the apical region of epithelial cells. To mimic this aspect *in vitro*, it is necessary to precisely control the inoculum amount, timing and cell targeting, which can be highly challenging in organoid systems ^18,19^. To address this, we employed an organ-on-chip technology to gain inoculation precision and reproducibility. When installed onto microfluidic chips, fetal-derived lung cells proliferated, differentiated and polarized in response to signaling cues present in the differentiation media. Multipotent organoid-derived cells differentiated on the airway chips and produced the expected goblet, club, basal, and multiciliated cell populations. The number of multiciliated cells derived on-chip was consistent with previous reports on primary cells cultured in this medium ^9^. Further optimization could include adding Notch inhibitors and BMP activators, as demonstrated by Van der Vaart et al. ^46^, to better control the ratio between the different cell types. In the case of alveolar chips, we observed a robust production of bona fide AT2 cells (SFTPC+, HT2-280+ and LAMP3+). This is in line with recent advancements for the generation of mature AT2 cells derived from fetal material ^6,26,47^. Previous efforts struggled with sustaining differentiated AT2 cells for more than few days in culture ^7^. Additionally, we observed a significant number of AT1 cells, detected with characteristic markers such as HT1-56, RAGE and CLIC5. AT1 cells were less abundant than AT2 cells in this model, with a 1:1.6 ratio based on HT1-56 and HT2-280 markers; however, this ratio is consistent with *in vivo* findings (1:2 or 1:3 ratio, as reported by Crapo et al. and Foster et al. ^48,49^). While increasing the AT1 population was not the focus of this study, it may be achievable on-chip by inducing AT2-to-AT1 transdifferentiation, as demonstrated in other studies ^14,29,43,48^. Overall, our characterization of the Alveochips validates their use for modeling airborne infections from the colonization steps to the innate immune response.

Previous studies have demonstrated the effects of *S. pneumoniae* infection in human immortalized cell lines ^35^, bronchial epithelial cultures ^50^ and alveolar organoids ^41^. Our work confirms the capacity of TIGR4 and 6B ST90 strains to colonize human alveolar epithelia. In murine models, the 6B ST90 strain does not cause disease whereas TIGR4 presents a highly invasive phenotype ^35^. Interestingly, in our model TIGR4 crosses the epithelial barrier to a greater extent than the 6B ST90 strain, partially phenocopying in vivo observations ^35,51^. In our experimental conditions, limited barrier damage was observed upon TIGR4 infection. While TIGR4 infection is commonly associated with epithelial cell death in most model systems ^34^, our AlveoChips experiments reveal a different dynamic. After 24hpi, high levels of TIGR4 counts were recovered from the chip bottom channel without a visible effect on the epithelial barrier, which was confirmed by cell counting and ZO-1 staining. These observations suggest that, at an early time point, TIGR4 can cross the epithelium without causing widespread damage. Altogether, our model provides a unique model for future investigations into how barrier level destruction may occur as disease progresses and whether recruitment of innate immune cells such as macrophages and neutrophils exacerbates this. Indeed, it is known that murine pneumococcal challenge with invasive disease favouring isolates leads to two waves of neutrophil infiltration within the lungs. Interestingly, the second wave coincides with rapid increase in bacterial burden and tissue damage, as opposed to the first being able to partially control pneumococci ^52^. On the other hand, we also found that *C. accolens*, a commensal nasal species, did not infect alveolar chips.–Previous studies have shown that *C. accolens* is highly abundant in the upper airways, where it can inhibit pneumococcal growth and colonization of the respiratory tract ^36,53^. Future work could explore whether co-culturing *C. accolens* and *S. pneumoniae* before chip inoculation affects pneumococcal infection of alveolar cells. The engineered AlveoChip system is capable of supporting investigations on pneumococcal transmigration events and host responsiveness within an alveolar epithelium, which are still poorly understood.

The COVID-19 pandemic showcased the continuous emergence of novel variants of concern, each using different mechanisms for viral spread and immune evasion to increase infectivity within the respiratory tract ^54,55^. SARS-CoV-2 effects have been widely investigated in airway and alveolar-like *in vitro* systems such as reconstructed airway epithelia in transwells ^5,30,56–58^, organoids ^10,39,57,59^ and organ-on-chip models ^45,60,61^. The general consensus is that novel variants, from the Omicron sublineage, replicate faster and more efficiently in the upper respiratory tract epithelium compared to earlier variants, which likely contributes to its increased transmissibility ^37,56,62^. In contrast, in *ex vivo* explant cultures of human lungs, Omicron replicates less efficiently than earlier variants such as Delta ^37^. Other studies using an immortalized alveolar cell line ^63^, iPSC-derived alveolar cells in micropatterned wells ^64^ and adult-derived alveolar organoids ^38^ also report the lower infectivity of Omicron in the alveolar microenvironment when compared to Delta. In this work we found that the AlveoChip model can recapitulate these replication differences between Delta and Omicron BA.5 variants.

Interestingly, we observed a marked increase in IFNα secretion following SARS-CoV-2 infection in AlveoChips. Both Delta and Omicron BA.5 triggered a robust release of IFNα, detectable at 4dpi. Our data also showed that IFN-β was also secreted mainly in Omicron BA.5-infected chips. The expression of ISGs following Delta and Omicron infection reflected the increased IFN secretion. Despite similar induction of the IFN response, the two variants displayed different replication efficiencies, suggesting that they vary in their ability to evade IFN signaling. In the airway microenvironment, Omicron is known to be highly replicative and appears to be resistant to IFN signaling. These characteristics seem to underlie its higher transmissibility and fitness ^62,65^. Recent single-cell RNA sequencing findings suggest that IFN resistance may be cell-type dependent ^57^. For instance, ciliated cells infected with SARS-CoV-2 elicit a clear IFN response, but secretory cells, which can also become infected, produce an even stronger IFN response, impeding viral replication at later stages ^57^. In pediatric nasal epithelial cells, goblet cells show a robust IFN expression, and consistently produce fewer viral particles than ciliated cells ^58^. Our findings that the strong IFN induction in alveolar cells exposed to Omicron viruses correlates with Omicron’s lower replication in this tissue suggest that SARS-CoV-2 exhibits cell- or tissue-specific IFN susceptibility.

In summary, our study presents a novel approach to building organ-on-chip models derived from human multipotent lung organoids. This strategy allowed the production of airway and alveolar faithful mimics. We showed that our model can recapitulate the early events following bacterial or viral infection in vitro. Our work also provides mechanistic understandings on early pathogenesis events and on human immune responses, that are otherwise inaccessible.

## Methods

### Procurement of human fetal lung tissue

Human fetal lung tissues were obtained from Advanced Bioscience Resources Inc. (CA, USA). Sourcing, import and organoid derivation were carried out as per national and institutional regulations and guidelines (import authorization number IE-2022-2353 and file number: 2022-075_Alveolar organoids at Institut Pasteur). Samples aged from 15 to 24 post-conception weeks. All the samples used for the current study had no known genetic abnormalities and were tested negative for Syphilis HIV, HBV, HCV, HSV, and CMV.

### Primary culture of human fetal lungs and establishment of organoid cultures

The procedure is based on protocols described by Nikolic et al. and Miller et al. ^24,25^. Briefly, distal regions of fetal lung lobes were cut into small 2 mm^3^ pieces in washing medium (100 units.mL^-1^ Penicillin, 100 µg.mL^-1^ Streptomycin and 0.25 µg.mL^-1^ Amphotericin B in HBSS). After several washes in the same medium, lung fragments were dissociated for 1h at 37 °C with intermittent agitation in a dissociation medium consisting of 1mg/ml Collagenase/Dispase (Roche), 40 U Dnase I (Thermo), 100 units.ml^-1^ Penicillin, 100 µg.ml^-1^ Streptomycin and 0.25 µg.mL^-1^ Amphotericin B in HBSS in RPMI base medium. The enzymatic reaction was inactivated by adding 10% Fetal Bovine Serum (FBS) and the samples were filtered through a 70 µm cell strainer, to filter out non-dissociated lung pieces. After centrifugation for 5min at 250 g, the samples were resuspended in a small volume of Advanced DMEM/F12 base medium with 100 units.ml^-1^ Penicillin, 100 µg.mL^-1^ Streptomycin and 1x Glutamax, then mixed with Geltrex (Thermo) in a 1:3 ratio. 40 µL drops were placed in 24 well plates and incubated at 37 °C for 30 min to solidify the gel. Finally, 400 µL of human self-renewal medium (hSRm) (Supplementary Table 1) were added to each drop and plates incubated under standard cell culture conditions (37 °C; 5% CO2). Medium was changed twice a week.

After 10-14 days of primary culture, lung organoids are ready for passage or cryopreservation. For passage, the medium was removed and 500 µL of cold Organoid Harvesting Solution (Cultrex) added to each well. Geltrex drops were removed, mechanically dissociated and collected in a 15 mL tube. Cold Advanced DMEM/F12 base medium was added up to 10 mL and the samples were centrifugated for 5 min at 100 g. The pellet was resuspended in undiluted Geltrex according to the desired splitting ratio (1:2 to 1:4). After gel solidification, 400 µL of hSRm medium was added in each well.

For freezing, Geltrex drops were detached and dissociated as described for passaging. After centrifugation, the pellets were resuspended in cold freezing medium (50% hSRm + 40% FBS + 10% DMSO), transferred overnight to the -80 °C freezer then to the liquid nitrogen for long term storage.

### Human fetal lung mesenchyme culture

During the first passaging of lung organoids, lung mesenchymal cells that had attached to the bottom of the well were recovered using 0.25% Trypsin-EDTA solution, followed by enzyme inactivation with 10% FBS in PBS. These cells were also retrieved after organoid centrifugation by collecting part of the supernatant above the organoid pellet. The cells were then incubated in 25 cm^2^ flasks coated with type I collagen (Sigma) and cultured in Advanced DMEM/F12 base medium supplemented with 10% FBS. Medium was changed twice a week until confluence. Lung mesenchymal cells were passaged several times and cryopreserved in the freezing medium detailed above.

### Airway and alveolar differentiation of lung organoids

Lung organoids were maintained in hSRm medium up to passage 5. Just after passaging this medium was replaced with either airway ^9^ or alveolar differentiation medium ^26^(Supplementary Table 1). Organoids were then cultured for an additional 2 weeks until fixation and immunostaining.

### Whole mount immunostaining of organoids

Organoids immunostaining was performed following the protocol described in Dekkers et al ^66^. Lung organoids were recovered from Geltrex drops and washed with cold DPBS (MgCl_2_+/CaCl_2_+) after centrifugation. Organoids were gently transferred to 1% bovine serum albumin (BSA) coated 15 mL tubes and fixed with 4% formaldehyde overnight at 4 °C. On the next day, the 15 mL tube was filled with 1% Tween in DPBS (MgCl_2_+/CaCl_2_+) (PBT) up to 10 mL, gently mixed, centrifugated at 70 g for 5 min at 4 °C. Organoids were either stored in PBT or used for immunolabeling.

For immunolabeling, organoids were first blocked for 1h in organoid washing buffer (OWB: 0.1% Triton + 0.2% BSA in DPBS) and then incubated overnight with primary antibodies using the same solution, with mild shaking at 4 °C. On the next day, the samples were washed with OWB for approximately 8 hours. The solution was changed every 2 hours. Secondary antibodies were incubated overnight in OWB with mild shaking at 4 °C. Samples were counterstained with 4′,6-diamidino-2-phenylindole (DAPI), transferred to imaging dishes (Ibidi) and imaged with a spinning disk confocal microscope Nikon Ti2E (Nikon). Images acquisition was performed with the NIS-Elements software (Nikon). The list of the primary and secondary antibodies used are shown in Supplementary Table 2.

### Lung on chip culture

Microfluidic S1^TM^ chips were purchased from Emulate Inc (Boston, USA). Chip handling and activation was performed as per the manufacturer’s indications. Briefly, one day before chip seeding, the devices were activated using ER1/ER2 reagents. Subsequently, top and bottom channels were incubated with 200 µg.mL^-1^ of human Collagen IV (C5533, Sigma) and 30 µg.mL^-1^ of Fibronectin (354008, Corning) at 37 °C overnight. The next day (day 0, see Figure 2A), the coating solution was removed and washed with DPBS, the chips were then ready for cell seeding.

Lung organoids up to passage 3 were recovered from Geltrex matrix as performed for passaging. The equivalent of 2 drops containing organoids were used for each chip. The organoids pellet was resuspended with an appropriate amount of hSRm medium (40 µL per chip) and mechanically fragmented using a P200 pipette. The level of fragmentation was verified by a standard benchtop microscope, to avoid the seeding of isolated cells. Chips were seeded with few microliters of cell suspension in the top channel and placed in an incubator at 37 °C overnight. The next day, hSRm medium was added to the cells. Non adherent cells were gently washed away. On day 2, sample-matched lung mesenchymal cells were detached from the culture flasks and seeded on the bottom channel at a density of 0.5 x 10^6^ cells per mL. In the meantime, Alveolar or Airway differentiation media (Supplementary Table 1) were added into top and bottom channel inlet reservoirs of the Pods (Emulate). When good cell attachment was reached, the chips were inserted into the Pods, placed into a ZOE CM1 device (Emulate) and top and bottom channels were perfused with medium at flow rate of 30 µL.h^-1^. On day 7, when the epithelial layer had reached 100% confluence, an air-liquid interface (ALI) was established by emptying media from the top channel. Thereafter, lung epithelial cells were fed through the bottom channel, perfused by the same differentiation medium. Chips were finally used for experiments on day 14 (alveolar chips) and day 21 (airway chips).

### Immunostaining on-chip

Chips were unmounted froms the Pods and fixed in formaldehyde 4% for 30 min, rinsed with DPBS and carefully cut in half using a scalpel blade. The samples were then placed in 24-well plates and treated with a permeabilization solution (0.1% Triton in DPBS) for 30 min, followed by blocking (2% BSA in DPBS) for 1 hour at room temperature. Next, primary antibodies diluted in 1% BSA in DPBS were incubated overnight at 4 °C. On the following day, secondary antibodies, diluted in 1% BSA in DPBS, were added for 2 hours. Samples were counterstained with DAPI and mounted in FluoromountG (Thermo Fisher Scientific) before observation with a spinning disk confocal microscope Nikon Ti2E (Nikon). Images were acquired with the NIS-Elements software (Nikon). Details of the primary and secondary antibodies and corresponding dilutions used are listed in Supplementary Table 2.

To quantify the percentage of positively stained cells, maximal intensity projections were generated using Fiji ^67^ and analyzed with QuPath version 0.5.1 ^68^ softwares. QuPath built-in tools were used for nuclei segmentation followed by cell classification using an interactive machine learning based approach. Percentage of positively stained cells was calculated as (number of stained cells/total nuclei number) x 100. Quantification of cell number per µm^2^ (Supplementary Figures 2A and 3A) was based on the ratio of counted nuclei per area of the chip.

### RNA extraction and quantitative real-time PCR

For RNA preparation, cells on chips were lysed with 350 µL of RNeasy Lysis buffer + 2% DTT (w/v). Total RNA was isolated using RNeasy microRNA kit according to the manufacturer’s instructions (Qiagen). RT-qPCR was performed in technical triplicates using the Luna Universal One-Step RT-qPCR Kit (New England Biolabs), following the manufacturer’s instructions. qPCR was performed on a QuantStudio 6 Flex thermocycler (Applied Biosystems, Thermo). Results are presented as relative RNA levels quantified by the 2-ΔΔCt method ^69^ and normalized to the endogenous control beta-actin. The gene expression profile was obtained from at least three independent experiments. Primer sequences are shown in Supplementary Table 3.

### Bacterial challenge of alveolar chips and bacterial enumeration

Inoculates of the Corynebacterium accolens (Ca) strain C781 were grown overnight in BHI broth supplemented with 1% Tween 80 ^70^. Experimental starter cultures of Streptococcus pneumoniae (Spn) isolates TIGR4 and 6B ST90 were prepared and grown as previously described in ref ^35^.

Briefly, stationary phase Ca or midlog Spn were collected by centrifugation, washed three times in DPBS followed by a 1:100 dilution in DPBS and dilution to 500 organisms per microliter using isolate specific CFU OD_600_ per mL conversion factors to make a challenge inoculum of 800 µL. Alveolar chips were challenged with 100 µl (50,000 organisms) of the challenge inoculum by perfusion through the apical input reservoir of the Emulate Pod at 1000 µL.h^-1^. Chips where exposed to the inoculum for 6 min.

Post-challenge inoculate was recovered from the output reservoir of the Emulate pod, pelleted, and suspended in 200 µL DPBS for CFU serial dilution plating (1:6). Bacterial adherence within the device was determined by subtracting the recovered post-challenge inoculate CFU counts from the counts of the challenge inoculum. Apical bacterial burden of the alveolar chips was determined by flushing the apical side with 500 µL of DPBS at 1000 µL.h^-1^ flow rate and collection of the perfused DPBS from the output reservoir. This was then pelleted and suspended in 200 µL DPBS for CFU serial dilution plating (1:6). Transmigrated bacteria across the alveolar barrier into the bottom chamber perfusion was determined by collecting total media from the bottom chamber output reservoir of the Emulate pod. This sample was pelleted by centrifugation and suspended in 200 µl DPBS for CFU serial dilution plating (1:6). All bacterial dilutions were plated on 5% Columbia blood agar plates and grown overnight at 37 °C with 5% CO_2_.

### SARS-CoV-2 infection of alveolar chips

Alveolar chips were cultured for 7 days in ALI conditions prior to SARS-CoV-2 infection. Chips and pods were then transferred from the ZOE CM1 device to a biosafety cabinet within a BSL3 laboratory. For each chip, 10^6^ RNA viral copies per µL of Delta and Omicron BA.5 viral stocks were used ^71,72^. Viral inputs were diluted in DMEM/F12 medium to a final volume of 200 μL and added into the pod’s top channel inlet reservoir. A 1 min pulse at a flow rate of 1000 μL.h^-1^ was programmed on the ZOE CM1 machine for viral inoculation. Chips were then incubated for 3 h at 37 °C, 5% CO2 under static conditions. Control chips were mock-treated with DMEM/F12 for the same duration. Post-infection, the remaining viral inoculum in the pods were collected for RT-qPCR analysis, and the chips were subjected to a series of washes. Briefly, 200 μL of DPBS was added to the top channel inlet reservoirs, followed by three 5 min pulses at 1000 μL.h^-1^ on a ZOE CM1 machine. In between each pulse we let the chips 5 min without perfusion, resulting in a total washing time of 30 min. A last pulse with DMEM/F12 was performed, followed by 20 min incubation at 37 °C. After this step, the media was collected in the outlet for RT-qPCR analysis. ALI conditions were then re-established. Apically released viruses were collected daily by loading 200 μL Advanced DMEM/F12 in the pod top channel inlet reservoir, followed by 1 min pulse at 1000 μL.h^-1^ in ZOE and by 20 min incubation at 37 °C. ALI was re-established after this step. On day 4, chips were either fixed in formaldehyde 4% for 30 min or the cellular RNA was isolated with RNeasy microRNA kit (Qiagen). Samples were stored until further analysis.

### Viral RNA quantification

Top and bottom channel effluents collected at days 0, 1 2 and 4 post-infection were inactivated for 20 min at 80 °C and subsequently subjected to RT-qPCR analysis with the Luna Universal One-Step RT-qPCR Kit (New England Biolabs), following the manufacturer’s instructions. SARS-CoV-2 RNA was quantified in a final volume of 5 μL per reaction in 384-well plates using SARS-CoV-2 N-specific primers on a QuantStudio 6 Flex thermocycler (Applied Biosystems, Thermo). Standard curve was performed in parallel using purified SARS-CoV-2 viral RNA ^5^.

### Scanning electron microscopy

Samples were fixed with 2.5% glutaraldehyde in 0.1 M cacodylate buffer for 1 h at room temperature and then for 12 h at 4 °C to inactivate SARS-CoV-2. Samples were then washed in 0.1 M cacodylate buffer and several times in water and incubated for 1 h at room temperature in 1% osmium tetraoxide. After dehydration by incubation in increasing concentrations of ethanol, samples were critical point dried with hexamethyldisilazabe, mounted on a stub, and analyzed by field emission scanning electron microscopy with a Jeol IT700HR microscope operating at 3 kV.

### Quantification of Interferons using ultra-sensitive digital ELISA (Simoa) assays

Prior to protein analysis, samples were treated in a BSL3 laboratory for viral inactivation as described by Robinot et al ^5^. Briefly, 2.5 µL of 10% Triton X-100 solution (Sigma) were added to 25 µL of each lung-on-chip apical lavage supernatant sample (1% final concentration) to be analyzed for IFNα, IFNβ and IFNλ3. Samples were then incubated 1 h at room temperature. Prior to Simoa quantification, inactivated samples were diluted 1:5 by adding 110 µL of Diluent B (Quanterix, Billerica, MA, USA) containing 11 µL of 10% Triton X-100 solution (1% final concentration). To estimate the limit of quantification, 8 blanks containing only detector/sample diluent were treated similarly.

IFNα, IFNβ and IFNl3 protein supernatant concentrations were quantified by Simoa digital ELISA assays developed with Quanterix Homebrew kits as previously described ^73–75^. After checking the absence of cross-reactivity, the three assays were multiplexed to decrease sample volumes needed. The SBG enzyme concentration was 150 pM.

For the IFNα assay, which quantifies all IFNα subtypes with a similar sensitivity, the 8H1 antibody clone was used as a capture antibody after coating on paramagnetic beads (0.3 mg.mL^-1^), the 12H5 clone was biotinylated (biotin/antibody ratio = 30/1) and used as the detector at a concentration of 0.3 µg.mL^-1^, and the 11150 recombinant human IFNα17 protein (PBL Assay Science, Piscataway, NJ, USA) was used as the standard. The limit of detection of the assay was 0.022 fg.mL^-1^ and the mean of the blanks was 0.46 fg.mL^-1^, including the dilution factor used (5.5).

For the IFNβ assay, the 710906-9 IgG1, kappa, mouse monoclonal antibody (PBL Assay Science) was used as a capture antibody after coating on paramagnetic beads (0.3 mg.mL^-1^), the 710323-9 IgG1, kappa, mouse monoclonal antibody (PBL Assay Science) was biotinylated (biotin/antibody ratio = 40/1) and used as the detector antibody at a concentration of 1 µg.mL^-1^, and the 11415 recombinant human IFNβ protein (PBL Assay Science) was used to quantify IFNβ concentrations. The limit of detection of the assay was 0.029 pg.mL^-1^ and the mean of the blanks was 0.17 pg.mL^-1^, including the dilution factor used (5.5).

For the IFNλ3 assay, the 21730-1 IgG1, kappa, mouse monoclonal antibody (PBL Assay Science) was used as a capture antibody after coating on paramagnetic beads (0.3 mg.mL^-1^), the MAB52591R IgG2a, mouse monoclonal antibody (R&D Systems, Minneapolis, MN, USA) was biotinylated (biotin/antibody ratio = 60/1) and used as the detector antibody at a concentration of 0.3 µg.mL^-1^, and the 5259-IL-025/CF recombinant human IFNλ3 protein (R&D Systems) was used as calibrator. The limit of detection of the assay was 0.006 pg.mL^-1^ and the mean of the blanks was 0.82 pg.mL^-1^, including the dilution factor used (5.5).

### Statistics and reproducibility

Statistical analyses were performed with the Prism software v8.4.3 (GraphPad). Unless stated otherwise, we used the nonparametric Mann–Whitney or Kruskal-Wallis tests. Statistical significance was assigned when p values were <0.05. The nature of statistical tests and the number of experiments (n) are reported in the figure legends.

## Acknowledgements

The authors would like to thank Olivier Schwartz and Florence Guivel-Benhassine for providing SARS-CoV-2 virus samples, Hugo Mouquet for providing anti-spike and anti-N antibodies and Sarah Vreugde for sharing bacterial strains. We thank the Single Cell Biomarkers UTechS of the Institut Pasteur for support in conducting this study. We acknowledge the Flow Cytometry Platform at Institut Pasteur for support in conducting this work. We thank the Biomics Platform (C2RT, Institut Pasteur) that is supported by France Génomique (ANR-10-INBS-09) and IBISA. We gratefully acknowledge the UtechS Photonic BioImaging, (C2RT, Institut Pasteur) supported by the French National Research Agency (France BioImaging; ANR-10–INBS–04; Investments for the Future). Experimental work benefited from the support of ultrastructure bioimaging core facilities of Institut Pasteur.

## Author contributions

BFF designed and conducted most of the experiments including organoid, organ-on-chip and SARS-CoV-2 infections; JWG derived lung bud organoid cultures and performed organoid differentiation experiments; MC carried out and analyzed bacterial infection experiments; HM and MHK assisted with organ-on-chip experiments; RY contributed to organoid and organ-on-chip experiments; VB performed Simoa analyses; VM performed electron microscopy; JDS and HSM contributed to the organoid derivation protocol; DD supervised the Simoa experiments; MH supervised the bacterial experiments, helped secure funding and edited the manuscript; NS assisted with the experimental designs and reviewed the data; LAC contributed to project conceptualization, SARS-CoV-2 infections and acquired funding for the project; SG conceptualized the project, secured funding, designed experiments and analyzed the data; BFF and SG wrote the manuscript. All authors read, edited and approved the final version.

---

**Supplementary Figure 1:**
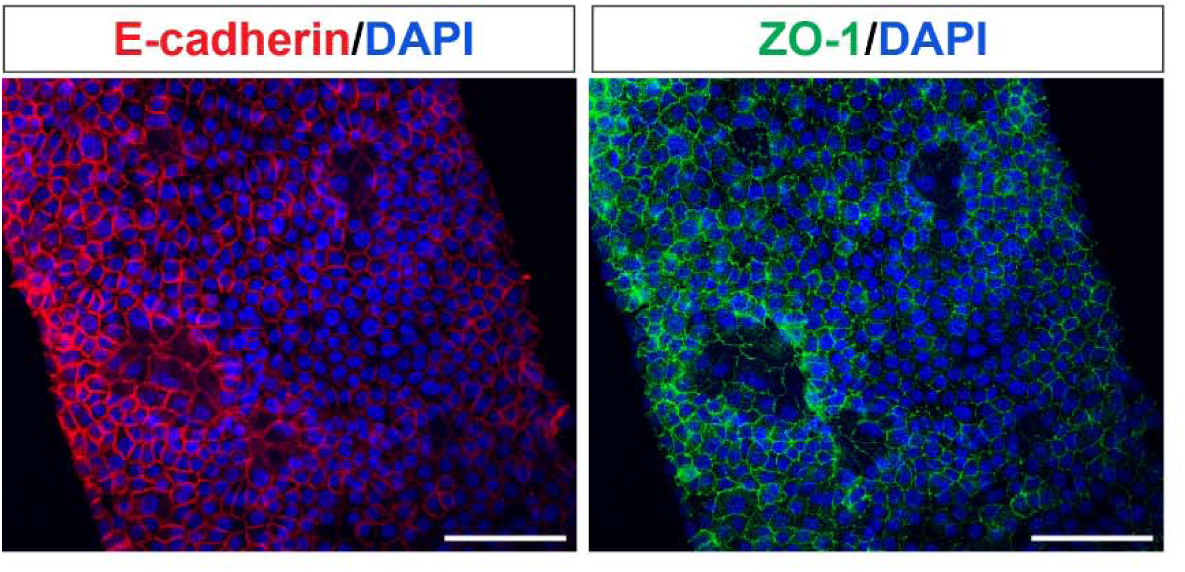
Multipotent lung organoids form a simple and tight epithelium on-chip. Cell adherent and tight junctions revealed with E-cadherin and ZO-1 immunolabeling, respectively. DAPI was used to counterstain nuclei. Scale bars: 100 µm.

**Supplementary Figure 2:**
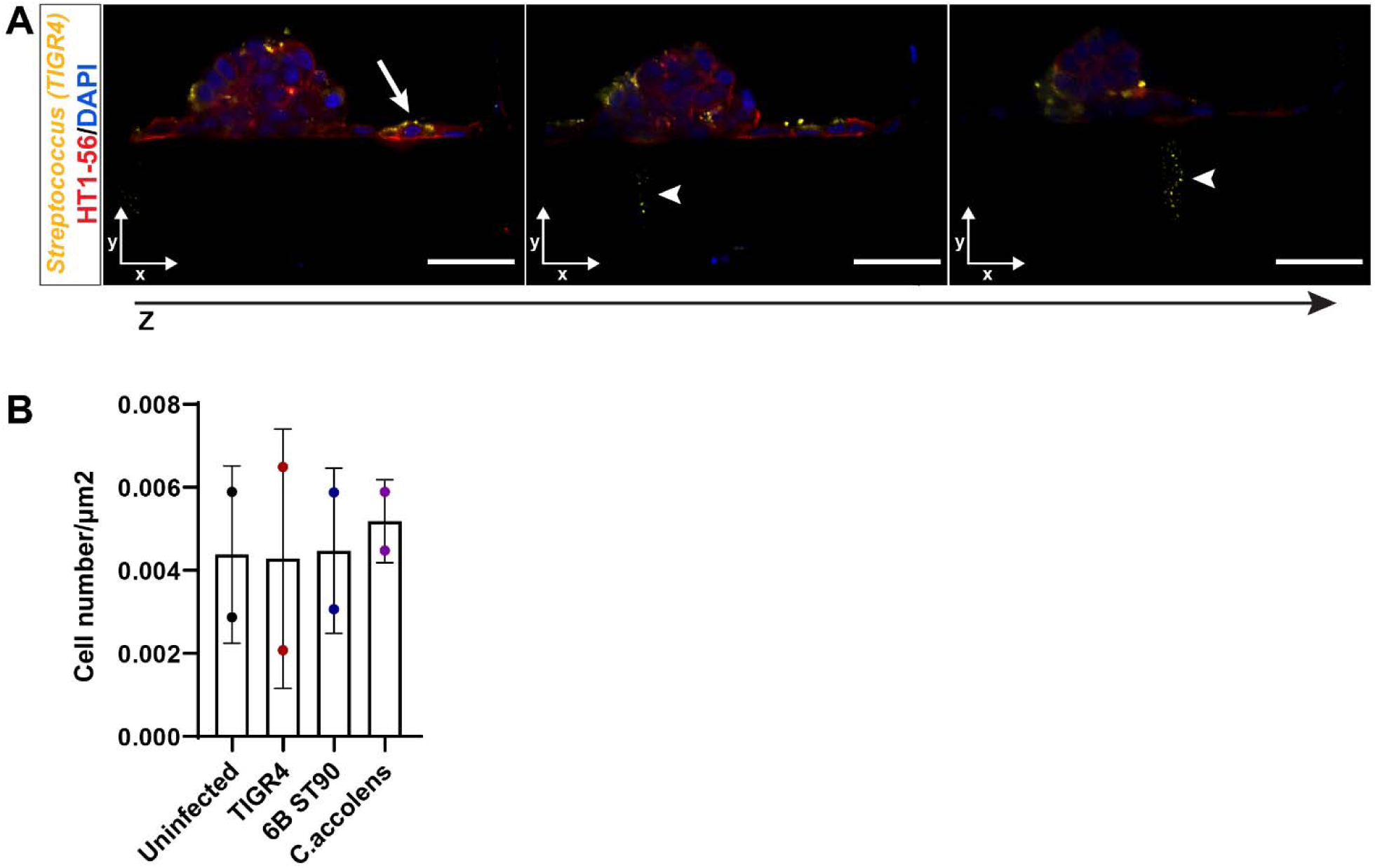
*Streptococcus pneumoniae* infection in human alveolar chips. (A) Quantification of number of cells per chip based on the ratio of nuclei number per um^2^ chip area. (B) Streptococcus immunolabeling of bacteria specimens surrounding the bottom channel lung mesenchyme at 24 hours postinfection (hpi). Scale bars: 100 µm.

**Supplementary Figure 3:**
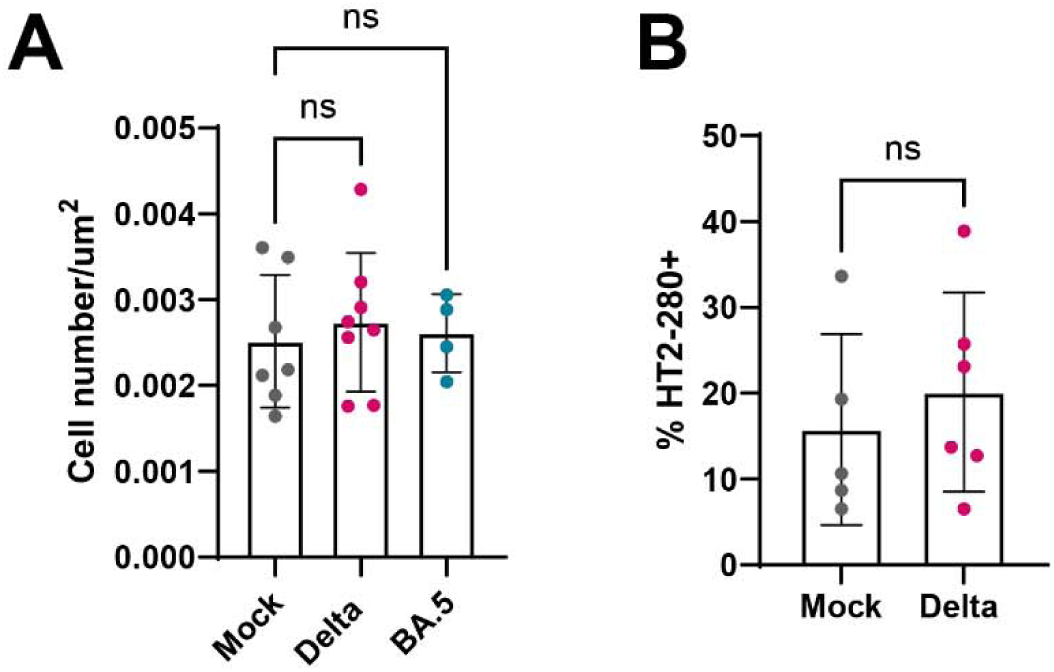
Effects of SARS-CoV-2 infection on alveolar cell populations. (A) Quantification of number of cells per chip based on the ratio of nuclei number per um^2^ chip area. (B) Percentage of total HT2-280+ cells per condition. Median +/- interquartile range are plotted and statistically significant differences are compared with the Kruskal-Wallis test.

**Supplementary Figure 4:**
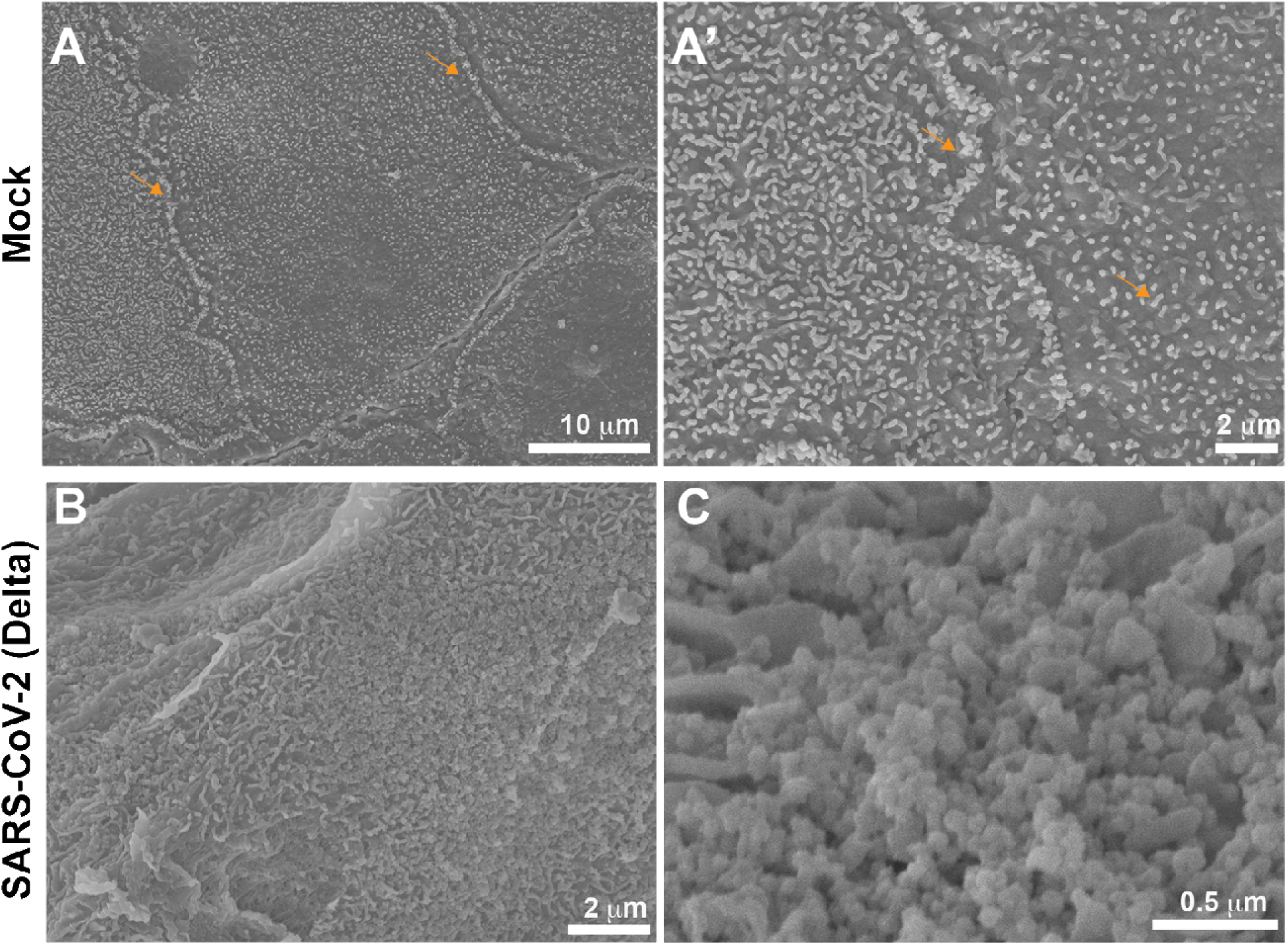
Ultrastructural features of AT2-like cells upon SARS-CoV-2 infection on-chip. (A) Scanning electron microscopy images evidence abundant small projections similar to microvilli in cells of an uninfected alveolar chip (arrows in A and A’). (B) Scanning electron microscopy images of a SARS-CoV-2 infected chip. (C) Viral particles observed at a higher resolution image than B.

**Supplementary Figure 5:**
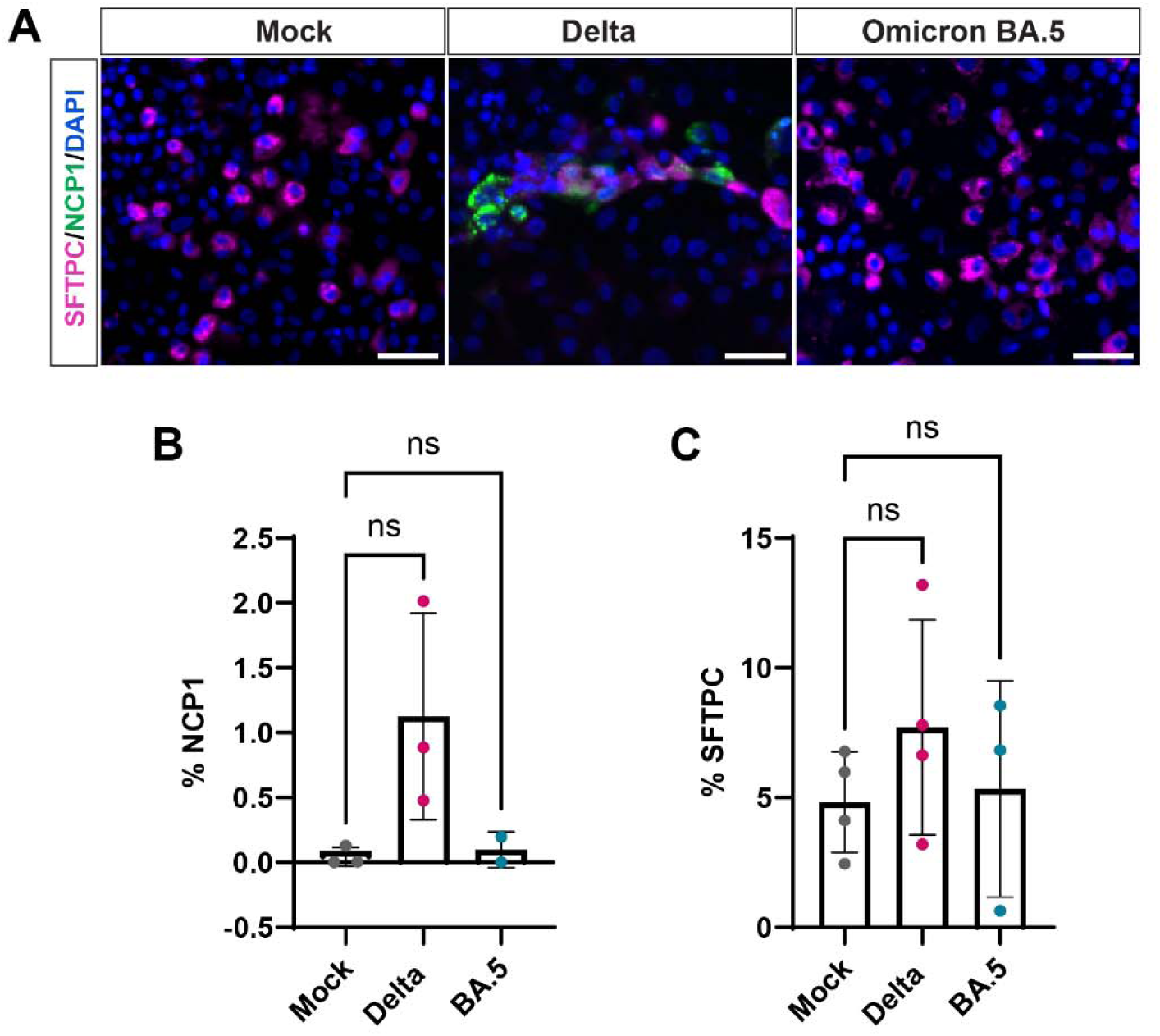
Comparison of SARS-CoV-2 variant infections in human alveolar chips. (A) Immunolabeling of SARS-CoV-2 infected cells with anti-NCP1-antibody (green). Alveolar cells are identified with an anti-pro-SFTPC antibody (magenta). DAPI is used to counterstain nuclei. Scale bars: 50 µm (B) Percentage of NCP1+ cells in different conditions at 4 dpi. (C) Percentage of pro-SFTPC+ cells in different conditions at 4 dpi. Median +/- interquartile range are plotted and statistically significant differences are compared with the Kruskal-Wallis test.

**Supplementary Figure 6.**
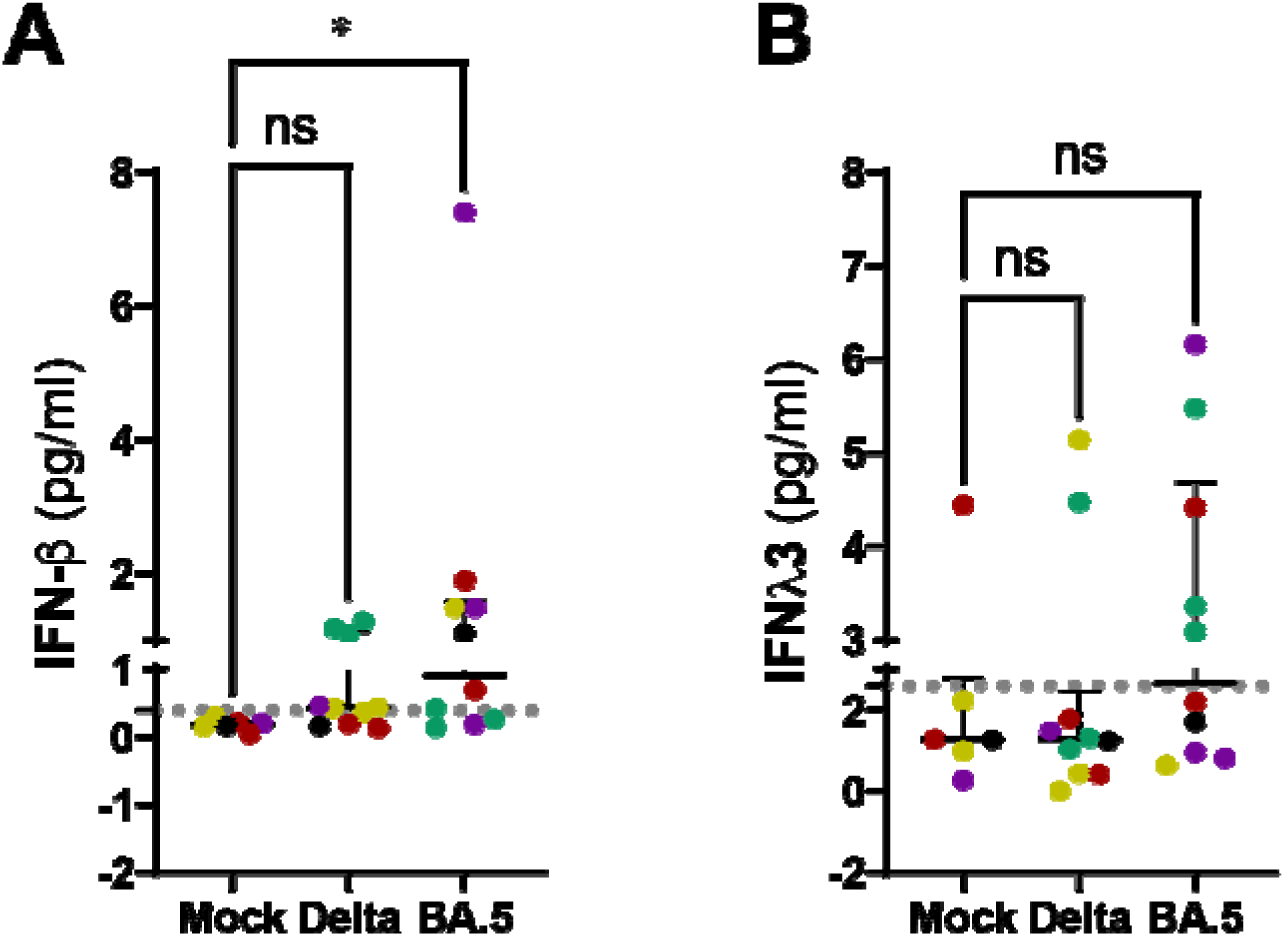
IFN host-response to SARS-CoV-2 infection in alveolar chips. **(A)** IFNβ and (B) IFNλ3 proteins detected in top channel washes recovered at 4 days post-infection. (n=5 independent experiments with 5 different organoid lines. Each dot color represents an organoid line. Median +/- interquartile range are plotted and Kruskal-Wallis tests are used for addressing statistically significant differences).

## References

1. Davis, J. D. & Wypych, T. P. Cellular and functional heterogeneity of the airway epithelium. Mucosal Immunology 14, 978–990 (2021).

2. Nikolić, M. Z., Sun, D. & Rawlins, E. L. Human lung development: recent progress and new challenges. Development 145, dev163485 (2018).

3. Mahieu, L., Van Moll, L., De Vooght, L., Delputte, P. & Cos, P. In vitro modelling of bacterial pneumonia: a comparative analysis of widely applied complex cell culture models. FEMS Microbiology Reviews 48, fuae007 (2024).

4. Huh, D. et al. Reconstituting Organ-Level Lung Functions on a Chip. Science 328, 1662– 1668 (2010).

5. Robinot, R. et al. SARS-CoV-2 infection induces the dedifferentiation of multiciliated cells and impairs mucociliary clearance. Nat Commun 12, 4354 (2021).

6. Sucre, J. M. S. et al. Successful Establishment of Primary Type II Alveolar Epithelium with 3D Organotypic Coculture. Am J Respir Cell Mol Biol 59, 158–166 (2018).

7. Dobbs, L. G. Isolation and culture of alveolar type II cells. American Journal of Physiology-Lung Cellular and Molecular Physiology 258, L134–L147 (1990).

8. Barkauskas, C. E. et al. Type 2 alveolar cells are stem cells in adult lung. J Clin Invest 123, 3025–3036 (2013).

9. Sachs, N. et al. Long-term expanding human airway organoids for disease modeling. The EMBO Journal 38, 1–20 (2019).

10. Salahudeen, A. A. et al. Progenitor identification and SARS-CoV-2 infection in human distal lung organoids. Nature 588, 670–675 (2020).

11. Dye, B. R. et al. In vitro generation of human pluripotent stem cell derived lung organoids. eLife 4, 1–25 (2015).

12. Yamamoto, Y. et al. Long-term expansion of alveolar stem cells derived from human iPS cells in organoids. Nature Methods 14, 1097–1106 (2017).

13. Jacob, A. et al. Differentiation of Human Pluripotent Stem Cells into Functional Lung Alveolar Epithelial Cells. Cell Stem Cell 21, 472–488.e10 (2017).

14. Burgess, C. L. et al. Generation of human alveolar epithelial type I cells from pluripotent stem cells. Cell Stem Cell 31, 657–675.e8 (2024).

15. Chen, Y.-W. et al. A three-dimensional model of human lung development and disease from pluripotent stem cells. Nature Cell Biology 19, 542–549 (2017).

16. Dichtl, S., Posch, W.* & Wilflingseder, D. The breathtaking world of human respiratory in vitro models: Investigating lung diseases and infections in 3D models, organoids, and lung-on-chip. European Journal of Immunology 54, 2250356 (2024).

17. Vazquez-Armendariz, A. I. & Tata, P. R. Recent advances in lung organoid development and applications in disease modeling. J Clin Invest 133, (2023).

18. Kim, M., Fevre, C., Lavina, M., Disson, O. & Lecuit, M. Live Imaging Reveals Listeria Hijacking of E-Cadherin Recycling as It Crosses the Intestinal Barrier. Current Biology 31, 1037–1047.e4 (2021).

19. Iakobachvili, N. et al. Mycobacteria–host interactions in human bronchiolar airway organoids. Molecular Microbiology 117, 682–692 (2022).

20. Youk, J. et al. Three-Dimensional Human Alveolar Stem Cell Culture Models Reveal Infection Response to SARS-CoV-2. Cell Stem Cell 27, 905–919.e10 (2020).

21. Walocha, R., Kim, M., Wong-Ng, J., Gobaa, S. & Sauvonnet, N. Organoids and organ-on-chip technology for investigating host-microorganism interactions. Microbes and Infection 105319 (2024) doi:10.1016/j.micinf.2024.105319.

22. Grassart, A. et al. Bioengineered Human Organ-on-Chip Reveals Intestinal Microenvironment and Mechanical Forces Impacting *Shigella* Infection. Cell Host & Microbe 26, 435–444.e4 (2019).

23. Stucki, A. O. et al. A lung-on-a-chip array with an integrated bio-inspired respiration mechanism. Lab Chip 15, 1302–1310 (2015).

24. Nikolić, M. Z. et al. Human embryonic lung epithelial tips are multipotent progenitors that can be expanded in vitro as long-term self-renewing organoids. eLife 6, 1–33 (2017).

25. Miller, A. J., et al. In Vitro Induction and In Vivo Engraftment of Lung Bud Tip Progenitor Cells Derived from Human Pluripotent Stem Cells. Stem Cell Reports 10, 101–119 (2018).

26. Lim, K. et al. Organoid modeling of human fetal lung alveolar development reveals mechanisms of cell fate patterning and neonatal respiratory disease. Cell Stem Cell 30, 20–37.e9 (2023).

27. Gonzalez, R. F., Allen, L., Gonzales, L., Ballard, P. L. & Dobbs, L. G. HTII-280, a biomarker specific to the apical plasma membrane of human lung alveolar type II cells. Journal of Histochemistry and Cytochemistry 58, 891–901 (2010).

28. van Riet, S. et al. Organoid-based expansion of patient-derived primary alveolar type 2 cells for establishment of alveolus epithelial Lung-Chip cultures. Am J Physiol Lung Cell Mol Physiol 322, L526–L538 (2022).

29. Lim, K. et al. A novel human fetal lung-derived alveolar organoid model reveals mechanisms of surfactant protein C maturation relevant to interstitial lung disease. The EMBO Journal 1– 26 (2025).

30. Lamers, M. M. et al. An organoid-derived bronchioalveolar model for SARS-CoV-2 infection of human alveolar type II-like cells. The EMBO Journal 40, 1–19 (2021).

31. Goltsis, O. et al. Influence of mesenchymal and biophysical components on distal lung organoid differentiation. Stem Cell Research & Therapy 15, 273 (2024).

32. Salaun, B. et al. CD208/dendritic cell-lysosomal associated membrane protein is a marker of normal and transformed type II pneumocytes. Am J Pathol 164, 861–871 (2004).

33. Treutlein, B. et al. Reconstructing lineage hierarchies of the distal lung epithelium using single-cell RNA-seq. Nature 509, 371–375 (2014).

34. Weiser, J. N., Ferreira, D. M. & Paton, J. C. Streptococcus pneumoniae: transmission, colonization and invasion. Nat Rev Microbiol 16, 355–367 (2018).

35. Connor, M. G. et al. The histone demethylase KDM6B fine-tunes the host response to Streptococcus pneumoniae. Nat Microbiol 6, 257–269 (2021).

36. Bomar, L., Brugger, S. D., Yost, B. H., Davies, S. S. & Lemon, K. P. Corynebacterium accolens Releases Antipneumococcal Free Fatty Acids from Human Nostril and Skin Surface Triacylglycerols. mBio 7, e01725–01715 (2016).

37. Hui, K. P. Y. et al. SARS-CoV-2 Omicron variant replication in human bronchus and lung ex vivo. Nature 603, 715–720 (2022).

38. Lamers, M. M. et al. SARS-CoV-2 Omicron efficiently infects human airway, but not alveolar epithelium. 2022.01.19.476898 Preprint at 10.1101/2022.01.19.476898 (2022).

39. Chiu, M. C. et al. A bipotential organoid model of respiratory epithelium recapitulates high infectivity of SARS-CoV-2 Omicron variant. Cell Discov 8, 1–15 (2022).

40. Huang, J. et al. SARS-CoV-2 Infection of Pluripotent Stem Cell-Derived Human Lung Alveolar Type 2 Cells Elicits a Rapid Epithelial-Intrinsic Inflammatory Response. Cell Stem Cell 27, 962–973.e7 (2020).

41. Sempere, J. et al. Minilungs from Human Embryonic Stem Cells to Study the Interaction of Streptococcus pneumoniae with the Respiratory Tract. Microbiology Spectrum 10, e00453–22 (2022).

42. Stucki, J. D. et al. Medium throughput breathing human primary cell alveolus-on-chip model. Sci Rep 8, 14359 (2018).

43. Bai, H. et al. Mechanical control of innate immune responses against viral infection revealed in a human lung alveolus chip. Nat Commun 13, 1928 (2022).

44. Barkal, L. J. et al. Microbial volatile communication in human organotypic lung models. Nat Commun 8, 1770 (2017).

45. Huang, D. et al. Reversed-engineered human alveolar lung-on-a-chip model. Proceedings of the National Academy of Sciences 118, e2016146118 (2021).

46. van der Vaart, J. et al. Modelling of primary ciliary dyskinesia using patient-derived airway organoids. EMBO reports 22, e52058 (2021).

47. Gonzales, L. W., Guttentag, S. H., Wade, K. C., Postle, A. D. & Ballard, P. L. Differentiation of human pulmonary type II cells in vitro by glucocorticoid plus cAMP. American Journal of Physiology-Lung Cellular and Molecular Physiology 283, L940–L951 (2002).

48. Foster, C. D., Varghese, L. S., Skalina, R. B., Gonzales, L. W. & Guttentag, S. H. In Vitro Transdifferentiation of Human Fetal Type II Cells Toward a Type I–like Cell. Pediatr Res 61, 404–409 (2007).

49. Crapo, J. D., Barry, B. E., Gehr, P., Bachofen, M. & Weibel, E. R. Cell number and cell characteristics of the normal human lung. Am Rev Respir Dis 126, 332–337 (1982).

50. Karwelat, D. et al. Influenza virus-mediated suppression of bronchial Chitinase-3-like 1 secretion promotes secondary pneumococcal infection. The FASEB Journal 34, 16432– 16448 (2020).

51. Sandgren, A. et al. Effect of Clonal and Serotype-Specific Properties on the Invasive Capacity of Streptococcus pneumoniae. The Journal of Infectious Diseases 189, 785–796 (2004).

52. Ghanem, E. N. B. et al. Extracellular Adenosine Protects against Streptococcus pneumoniae Lung Infection by Regulating Pulmonary Neutrophil Recruitment. PLOS Pathogens 11, e1005126 (2015).

53. Horn, K. J. et al. Corynebacterium Species Inhibit Streptococcus pneumoniae Colonization and Infection of the Mouse Airway. Front Microbiol 12, 804935 (2021).

54. Lamers, M. M. & Haagmans, B. L. SARS-CoV-2 pathogenesis. Nat Rev Microbiol 20, 270– 284 (2022).

55. Zhang, Y., Chen, S., Tian, Y. & Fu, X. Host factors of SARS-CoV-2 in infection, pathogenesis, and long-term effects. Front Cell Infect Microbiol 14, 1407261 (2024).

56. Planas, D. et al. Distinct evolution of SARS-CoV-2 Omicron XBB and BA.2.86/JN.1 lineages combining increased fitness and antibody evasion. Nat Commun 15, 2254 (2024).

57. Meganck, R. M. et al. SARS-CoV-2 variant of concern fitness and adaptation in primary human airway epithelia. Cell Reports 43, 114076 (2024).

58. Woodall, M. N. J. et al. Age-specific nasal epithelial responses to SARS-CoV-2 infection. Nat Microbiol 9, 1293–1311 (2024).

59. Katsura, H. et al. Human Lung Stem Cell-Based Alveolospheres Provide Insights into SARS-CoV-2-Mediated Interferon Responses and Pneumocyte Dysfunction. Cell Stem Cell 27, 890–904.e8 (2020).

60. Thacker, V. V. et al. Rapid endotheliitis and vascular damage characterize SARS-CoV-2 infection in a human lung-on-chip model. EMBO Rep 22, e52744 (2021).

61. Domizio, J. D. et al. The cGAS-STING pathway drives type I IFN immunopathology in COVID-19. Nature 145–151 (2022).

62. Shi, G. et al. Omicron Spike confers enhanced infectivity and interferon resistance to SARS-CoV-2 in human nasal tissue. Nat Commun 15, 889 (2024).

63. Mache, C. et al. SARS-CoV-2 Omicron variant is attenuated for replication in a polarized human lung epithelial cell model. Commun Biol 5, 1–8 (2022).

64. Masui, A. et al. Micro-patterned culture of iPSC-derived alveolar and airway cells distinguishes SARS-CoV-2 variants. Stem Cell Reports 1–17 (2024).

65. Otter, C. J. et al. Interferon signaling in the nasal epithelium distinguishes among lethal and common cold coronaviruses and mediates viral clearance. Proceedings of the National Academy of Sciences 121, e2402540121 (2024).

66. Dekkers, J. F. et al. High-resolution 3D imaging of fixed and cleared organoids. Nat Protoc 14, 1756–1771 (2019).

67. Schindelin, J. et al. Fiji: an open-source platform for biological-image analysis. Nature Methods 2012 9:7 9, 676–682 (2012).

68. Bankhead, P. et al. QuPath: Open source software for digital pathology image analysis. Sci Rep 7, 16878 (2017).

69. Livak, K. J. & Schmittgen, T. D. Analysis of Relative Gene Expression Data Using Real-Time Quantitative PCR and the 2−ΔΔCT Method. Methods 25, 402–408 (2001).

70. Menberu, M. A., et al. *In vitro* and *in vivo* evaluation of probiotic properties of *Corynebacterium accolens* isolated from the human nasal cavity. Microbiological Research 255, 126927 (2022).

71. Planas, D. et al. Reduced sensitivity of SARS-CoV-2 variant Delta to antibody neutralization. Nature 596, 276–280 (2021).

72. Bruel, T. et al. Longitudinal analysis of serum neutralization of SARS-CoV-2 Omicron BA.2, BA.4, and BA.5 in patients receiving monoclonal antibodies. Cell Reports Medicine 3, 100850 (2022).

73. Smith, N. et al. Distinct systemic and mucosal immune responses during acute SARS-CoV-2 infection. Nat Immunol 22, 1428–1439 (2021).

74. Llibre, A. et al. Plasma Type I IFN Protein Concentrations in Human Tuberculosis. Front Cell Infect Microbiol 9, 296 (2019).

75. Rodero, M. P. et al. Detection of interferon alpha protein reveals differential levels and cellular sources in disease. J Exp Med 214, 1547–1555 (2017).

